# Bridging the resolution-sectioning gap in stimulated emission depletion microscopy: theory and experiment

**DOI:** 10.1101/2024.01.18.576258

**Authors:** Michael Belsley, Joana Soares-de-Oliveira, António J. Pereira

## Abstract

Microscopes generally exhibit superior performance in 2D compared to 3D. Fluorescence super-resolution imaging often intensifies this discrepancy, such as with the gold-standard vortex-based stimulated emission depletion (STED) microscope, which narrows the point-spread function only laterally. In this study, we developed a semi-analytical theory based on the Nijboer-Zernike expansion, conducted simulations and performed experiments to establish the merits of the alternative bivortex-based STED. We find that this mode reduces the axial-lateral resolution mismatch and effectively emulates noisier multi-beam approaches by providing access to point-spread function geometries that are strictly forbidden to the two conventional single-beam modes, 2D-STED and z-STED. Notably, theory and experiment indicate that, besides filling the gap, bivortex STED delivers a higher signal-to-background than the two modes it bridges.

## Introduction

Stimulated emission depletion (STED) microscopy was proposed ^1^ and implemented ^2^ at the end of last century, becoming the first ever far-field super-resolution fluorescence microscopy technique. As with virtually all super-resolution techniques, the crux lies in the nonlinear interaction with a sample. STED uses non-linearity to write modulations at the sub-diffraction level, specifically by saturating fluorescence quenching around a point of interest that remains unquenched, which can, in principle, be made arbitrarily small. Despite the high optical fluxes necessary for saturation, its ease of use, speed, and minimal image artifacts have led to its wide adoption in bio-imaging. This acceptance increased significantly after the use of a helical phase ramp was suggested^3^ in 2004. This phase mask, commonly referred to as a vortex mask or spiral phase plate, is now considered an optimal solution for the generation of the required doughnut depletion beam. Along with maximal lateral steepness^3, 4^, the crucial intensity-null formed at the center of this beam survives common perturbations such as spherical aberration, objective pupil under-filling by the depletion laser or, more generally, any complex amplitude perturbation with cylindrical symmetry ^5-9^. While these perturbations will deteriorate the overall depletion beam shape, their impact is limited because it is really the darkness and the steepness of the intensity well that dominate performance^10^. Because of its robustness and maximal lateral resolution, the vortex-based implementation, or 2D-STED, is by far the most commonly used mode.

The depletion doughnut in 2D-STED does not, however, improve on the confocal microscope’s axial performance ^11, 12^. In particular, the capacity to filter out DC background and resolve different planes – a property termed optical sectioning^13^ - remains at the confocal level. 2D-STED thus aggravates the lateral-axial mismatch, which, together with ever more challenging signal conditions as saturation increases, presents a drawback in thicker and noisier biological specimens, where light arising from fluorophores outside of the focal plane often reduce contrast significantly.

To surpass confocal-level optical sectioning, the resort has been z-STED, which was actually the original embodiment of any far-field super-resolution microscope^2^. There, a top-hat phase mask creates two strong depletion lobes along the optical axis ^14-16^, just above and just below the focal plane, creating a photo-physical ‘pinhole’ at the sample. By working in series with the confocal pinhole, the z-STED pinhole will deliver super-confocality. Unfortunately, this mode suffers from well-known shortcomings^9, 17, 18^. One is that in order to achieve the intensity-null at the center of the doughnut, the mask’s radial scale (top-hat radius) must be calibrated for instrumental variables such as the refractive index profile between objective and object (e.g. oil-glass-water) and the objective’s fill factor of the incident laser beam. Although proper calibration and adaptive optics techniques ^19-21^ may be used for beam optimization, two problems remain that are inherent to the z-STED beam geometry and therefore hold true for a perfect beam. Namely, a comparatively modest lateral super-resolution ^2, 22^ and an incomplete coverage of the excitation beam’s secondary peaks, which will thus fluoresce, creating the so-called ghost-images ^22-24^.

Being a result of resolution-centric heuristics and optimizations^4^, 2D-STED and z-STED modes are ultimately distant and actually discrete points in the two-dimensional space of lateral-axial performance. To fill the gap, these depletion beams can be superimposed incoherently – a regime known as 3D-STED ^9, 25-27^, but the z-STED component’s own sensitivity to radial aberrations remains present in the 3D-STED compound beam and clearly can’t be cancelled via incoherent mixtures. Notably, even ideal component beams will generally not create an ideal compound beam due to the inevitable misalignments that result in an elevation of the doughnut’s ‘zero’, which will inadvertently and disproportionally^10^ deplete the often dim fluorescence signal. An alternative to 3D-STED that uses a single depletion beam was recently proposed^24^ that avoids beam misalignments and minimizes the impact of aberrations. Called coherent-hybrid STED (CH-STED), the depletion beam is formed by coherently adding a vortex-diffracted field to that generated by a free-scale top-hat, effectively forming a bivortex phase mask. On its own, such top-hat component (in general, a highly defective z-STED mask) couldn’t possibly create an on-axis intensity-zero because the inner and outer ‘beams’ are intentionally power-unbalanced, a situation where full destructive interference is infeasible. However, added to a vortex, it will always create an intensity-zero and, crucially, it is aberration-resilient^28^. Most important is the fact that this coherent-hybrid beam generated by the bivortex is shaped as a smooth concave capsule (bottle beam), even if incompletely closed because of a persistent dark node along the optical axis.

The key point that allows axial confinement is that the bivortex (CH-STED), but not the vortex (2D-STED), contains off-axis phase gradients, a necessary condition to get axial intensity modulations. That this is a requirement, hence fulfilled in z-STED also, can be elucidated by drawing an analogy to the hyperbolic fringes in the Young’s double-slit experiment, where the axial intensity profile will only display modulations (the ingredient for confinement) if the profiled line is offset from the optical axis. In practice, by varying the distance between the phase step and the optical axis, the beam undergoes a transformation that has heuristically placed CH-STED as interpolating over the gap between 2D-STED and z-STED-like behavior in imaging^24^ and spectroscopy, such as STED-FLIM^28^. The definition of a meaningful metric space where the actual location and interpolation span of CH-STED can be characterized would support STED users in making an informed choice of which mode to choose in each experimental situation.

In this work, we quantitatively explore the merits and drawbacks of CH-STED regarding lateral and axial performances and signal-to-background conditions, always using the standard STED modes as benchmarks. We approach these objectives through experiments with synthetic and biological specimens, use of a full vectorial diffraction simulation package ^29^ as well as a more insightful semi-analytical pseudo-paraxial scalar development based on a Nijboer-Zernike aberration expansion, which proves reliable in characterizing the point-spread function geometry of the system at scales well beyond the wavelength, a necessary and mostly sufficient condition to describe the super-resolution STED microscope.

## Results

### Depletion beam geometry: theory

To arrive at a semi-analytical description for the geometry of different depletion beams used in STED microscopy, namely 2D, z and CH, we followed the Nijboer-Zernike expansion developed for aberration analysis ^30-32^ (detailed in the **Supplemental Material**). To simplify the analysis, we employ a scalar approximation and assume that the depletion beam overfills the pupil of the objective lens so that plane waves of amplitude *A* are incident on the objective’s entrance pupil. Given these assumptions, the field amplitude at a point 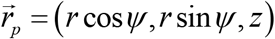 near the on-axis focal plane can be estimated through the Debye-Wolf diffraction integral

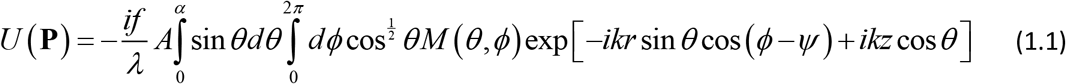

Here the spherical coordinates (*θ,ϕ*) define a unit vector pointing from an arbitrary point in the exit pupil towards the geometric focus of the objective, while k is the wavevector in image space. For an aplanatic system free from aberrations the amplitude strength factor is 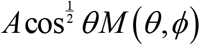 where *M* (*θ,ϕ*)determines the type of STED beam, describing the phase variation of the plane waves incident on the entrance pupil of the infinity corrected objective. In effect, the function *M* (*θ,ϕ*)can in principle be a complex amplitude, a generalization that relaxes the plane-wave approximation and can accommodate the Gaussian beam truncation that is usually found in practice, provided the beam is centered on the optical axis. The angle *α* is the maximum inclination of a ray within the objective’s exit pupil towards the geometric focus, with the numerical aperture *NA* = *n* sin *α, n* being the refractive index in image space.

As detailed in the **Supplemental Material** it is convenient to use scaled axial and radial coordinates

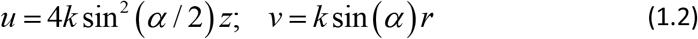

Then, making the coordinate transformation to a normalized exit pupil radius *σ* = sin θ/ sin *α* and neglecting a multiplicative factor in the integrand of 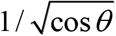, the Debye-Wolf integral becomes

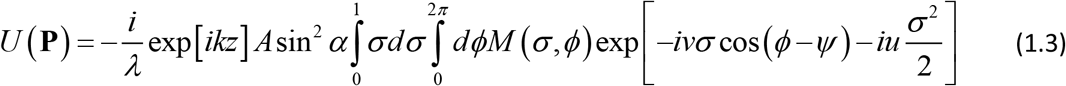

We have followed the recommendation of Sheppard and Matthews ^33^ and made a pseudo-paraxial approximation for the cos *θ* factor in the exponential,

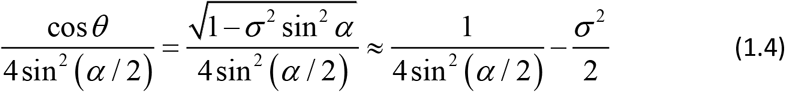

This approximation matches the limiting values of the phase term on-axis and at full aperture, and is often used to obtain improved estimates over the usual paraxial approximation. The conventional paraxial approximation expands the square root in the above expression in the small aperture limit.

The phase mask *M* (*ρ,θ*) in Eq.(1.3) determines the type of beam created. All STED beams studied in this work can be described by writing the phase mask in the general form,

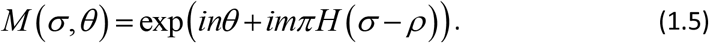

Here *H* is the Heaviside function and *ρ* denotes the scaled radius at which a step of *m π*radians occurs. This expression can describe a flat mask (e.g. to generate the excitation beam) (*n* = 0, *m* = 0), a z-STED mask (*n* = 0, *m* =1, *ρ*_*step*_ ≅ <2^−12^), a 2D-STED vortex mask (*n* = 1, *m* = 0) and a CH-STED bivortex mask (*n* = 1, *m* = 1(*typ*.), 0 < *ρ*_*step*_ < 1).

The case of a flat phase mask, *M* (*σ, θ*) = 1, provides a reference for the signal strength of undepleted emitters. As detailed in the **Supplemental Material**, using the Nijboer expansion, one quickly arrives at the following expression for the field amplitude,

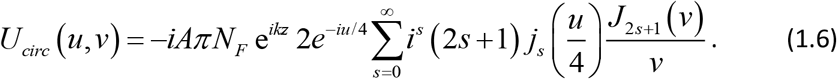

with *N*_*F*_ the Fresnel number for the objective. This series converges quickly and the first four terms are sufficient to obtain an intensity profile accurate to a few percent for (*u, v*) ranging from 0 to ± 10*π*. In a typical high-resolution fluorescence microscope (*NA*=1.4, *n*=1.5) this amounts to a range larger than ±2*λ*. However, we note that even a minor error in field estimation can be magnified when integrating azimuthally around the optical axis, as we will need to do below to assess the signal-to-background conditions.

To construct the z-STED depletion beam subtract twice the field diffracted by the inner disk phase region from that diffracted by the full flat phase mask

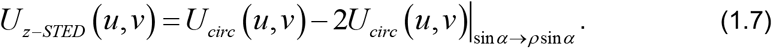

will create a zero-intensity central node only if the radius of the top hat, *ρ*, is chosen to divide the entrance pupil into two concentric regions of the same area (for a plane wave incident in a paraxial system), but careful tuning of *ρ* is required for high aperture systems or finite width beams (**Methods**). The transformation of *u, v* and *U*_00_ when sin*α → ρ* sin*α* is delineated in Eq S.15 of the **Supplemental Material**.

For the case of the single-vortex phase mask, as in 2D-STED, the field near the focal plane is given in the Nijboer-Zernike expansion as

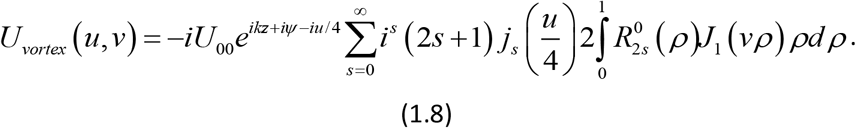

Here *U*_00_ = −*iAπN*_*F*_ is the focal plane field amplitude on axis for the case of a flat phase mask, corresponding to the peak field of the Airy spot. 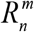 are the radial Zernike polynomials of order *n*.

Being scalar, the above treatment does not account for polarization, an aspect that becomes increasingly relevant in high-numerical aperture systems^34, 35^. Heuristically, the non-negligible axial components of the electric field may be cancelled by employing a circularly polarized beam, so that at least the nodal points can be recapitulated under a paraxial approach. A more comprehensive understanding of the match between the Nijboer-Zernike and vectorial simulations (and experiments) will be reached in the next sections. From symmetry arguments it is also easy to see that any radial perturbation (phase or amplitude) can be integrated to the nominal mask function (*σ, θ*)without compromising the on-axis interference conditions between all pairs of diametrical points at the pupil. Such perturbations can be radial aberrations (e.g., spherical) and/or radial attenuation (e.g., an under-filling Gaussian beam). This robustness of the nodal line along the optical axis is one reason that the single vortex 2D-STED has become the standard of STED microscopy ^3, 11, 36^.

For CH-STED the depletion beam is created by a bivortex ^24^, which can be viewed as a vortex to which a π-phase disk is added (Figure 1a). Since the phase disk is a strictly radial perturbation to the vortex, all the arguments just made regarding robustness of the nodal line to aberrations also apply to the bivortex (or multi-vortex). To retrieve the actual beam structure we proceed similarly to the top-hat case, now with *U*_*circ*_ replaced by *U*_*vortex*_. If we allow also for an arbitrary phase disc height *ϕ*_*step*_, the field amplitude becomes

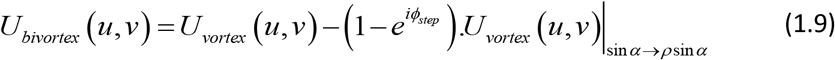

**Figure 1.**
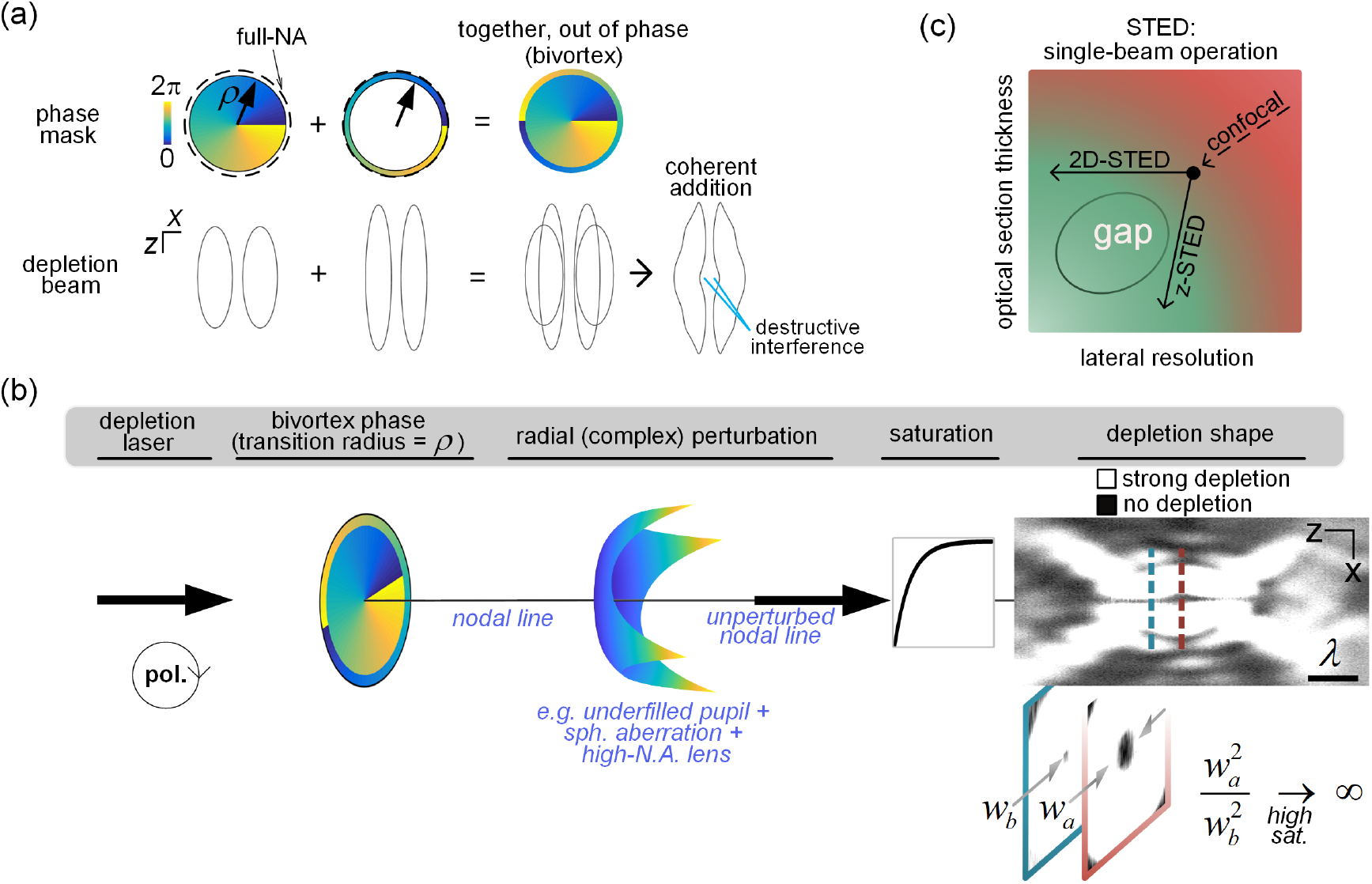
CH-STED overview. (a) Depletion beam construction via the vortex disc + vortex ring interpretation. The elementary fields assigned to these regions interfere destructively near focus. (b) A circularly polarized beam with cylindrically symmetric complex amplitude (e.g. a truncated Gaussian beam) is modulated by a bivortex mask and by a combination of radial perturbations, which includes the focusing lens. The optical axis is dark with a weak-depletion capsule formed in the focus region. The beam was ‘imaged’ by detection of the scattering signal from a gold nano-bead. The image is shown digitally clipped to convey the notion of saturation - the key ingredient in rendering the weight of the defocused portions of the peri-nodal line negligible (*w*_*b*_<<*w*_*a*_). (c) When using 2D-STED or z-STED, the relative weight of axial and lateral confinement cannot be chosen, leaving a gap in the performance space of single-beam STED setups.

Since *U*_*vortex*_ vanishes along the optical axis, it becomes clear from Eq. (1.9) that *U*_*bivortex*_ also does, regardless of the values of *ρ* and *ϕ*_*step*_. This is in stark contrast with the z-STED field, *U*_*top* −*hat*_ in Eq.(1.7), whose terms ∝*U*_*circ*_ are finite and therefore cancel only under certain conditions, namely for specific (*ρ,ϕ*_*step*_). As we will see, if *v ≠*0 the second term on the r.h.s. will lead to an inversion of intensity concavity sign if a threshold is crossed in the space (*ρ,ϕ*_*step*_), as required for axial confinement. The bivortex-diffracted depletion beam is the core of the CH-STED mode.

### Depletion beam geometry: experiment versus theory

The robustness of the intensity-zero at the center of the depletion beam is warranted in CH-STED and in 2D-STED as long as non-cylindrical complex perturbations are avoided. The downside of this robustness is that it implies that defocus (quadratic perturbation term) is also ‘allowed’, so that in both cases the intensity-null is not a point but a nodal line all along the optical axis. It was precisely this original shortcoming of the 2D-STED beam, which motivated the replacement of the vortex by a bivortex ^24^. Specifically, the concavity generated in the latter renders the weight of the peri-nodal ‘valley’ asymptotically negligible by virtue of saturation (Figure 1b). This means that, right after the action of the nano-second depletion beam, the off-focus planes carry even less integrated excited-state energy relative to the focal plane than in a confocal microscope, which is the definition of optical sectioning.

Sectioning improvement in CH-STED may at least partially fill the resolution-sectioning space that is inaccessible using previous single-beam STED strategies (Figure 1c).

To explore this possibility, we first characterize the shape of the depletion beam and later its effect on the fluorescence PSF of the microscope. To obtain depletion beam profiles experimentally, xz-scans were run to collect the signal scattered by sub-diffraction sized gold beads, without using a confocal pinhole. In Figure 2 we show the intensity cross-sections along a plane containing the optical axis for flat-phase and modulated depletion beams as obtained experimentally, via the Nijboer-Zernike expansion and also using a vectorial diffraction simulation package, as described in Leutenegger et al.^29^. To facilitate the comparison between STED modes, the Airy spot maximum *I* _*Airy*_, as obtained with the depletion laser hitting a flat-phase mask, is used as a global reference intensity for all depletion beam types.

**Figure 2.**
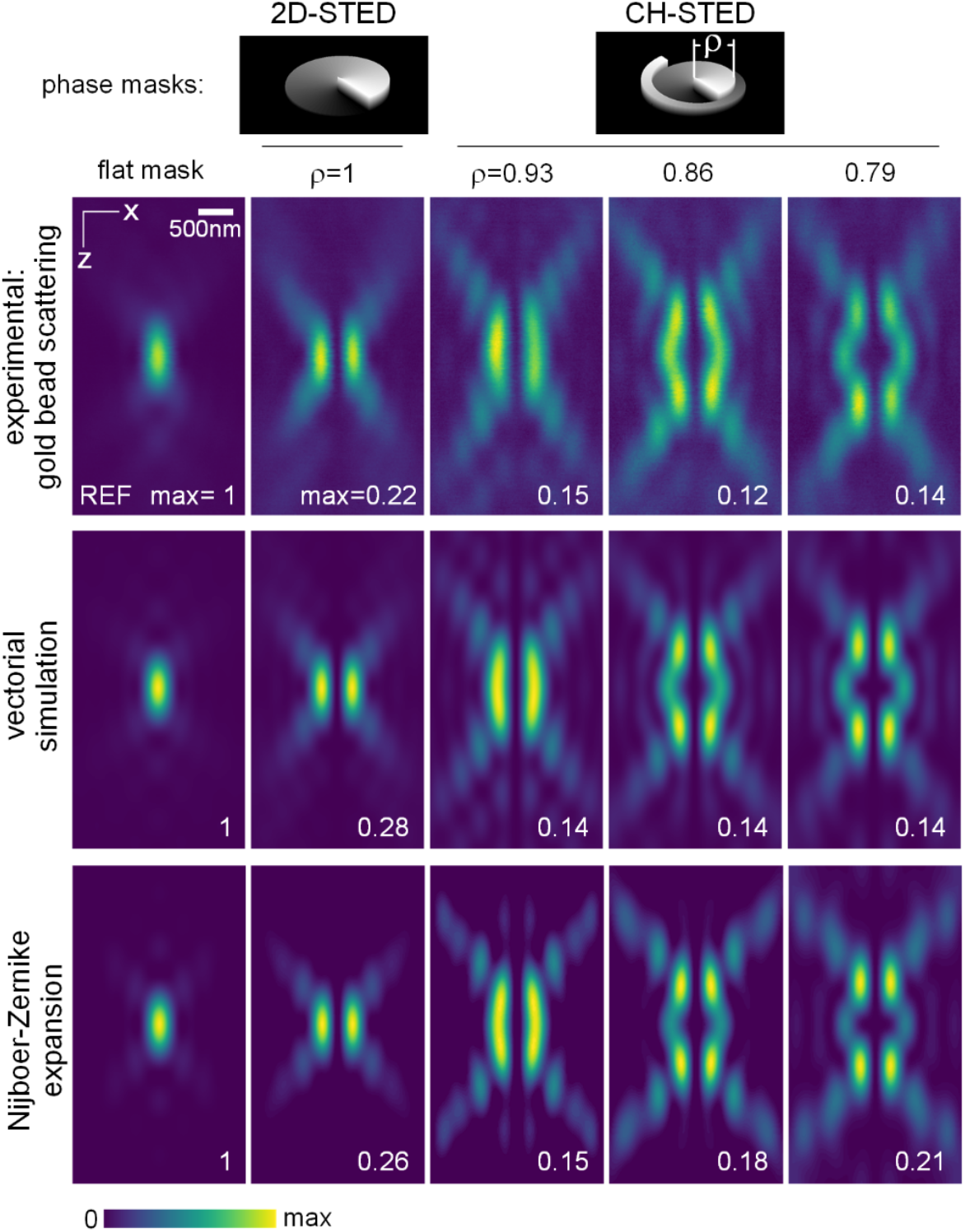
Depletion beam cross-sections from experiment and theory. Experimental, theoretical and simulated beam cross-sections are shown for the cases of a flat reference phase mask (maximum intensity=1) and several bivortex radii (ρ), including the particular case of 2D-STED (ρ=1). The experimental depletion beam is generated by applying the modulation to an SLM and observed by measuring the signal scattered from gold nano-beads. λ=775nm, *n*=1.52, N.A.=1.4.

### Photo-physical pinhole threshold

Under appropriate conditions, the beam generated by a bivortex, but not by a vortex (ρ=1), will form a photo-physical pinhole inside the sample, defining a qualitative change essential for STED confinement. To determine the bivortex radius threshold at which this transition occurs, we calculated the axial curvature across the focal plane as a function of ρ and of the distance from the optical axis (Figure 3a). The concavity changes sign at a bivortex radius ρ=0.94-0.95. Except at the optical axis, where the axial concavity is always zero, this threshold remains essentially constant as the measurement profile is displaced away from the optical axis within the λ*⁄*(2*NA*) range, so the isophote surfaces crossing the focal plane switch curvature synchronously, supporting the concept of a global pinhole threshold (Figures 3b and S4).

**Figure 3.**
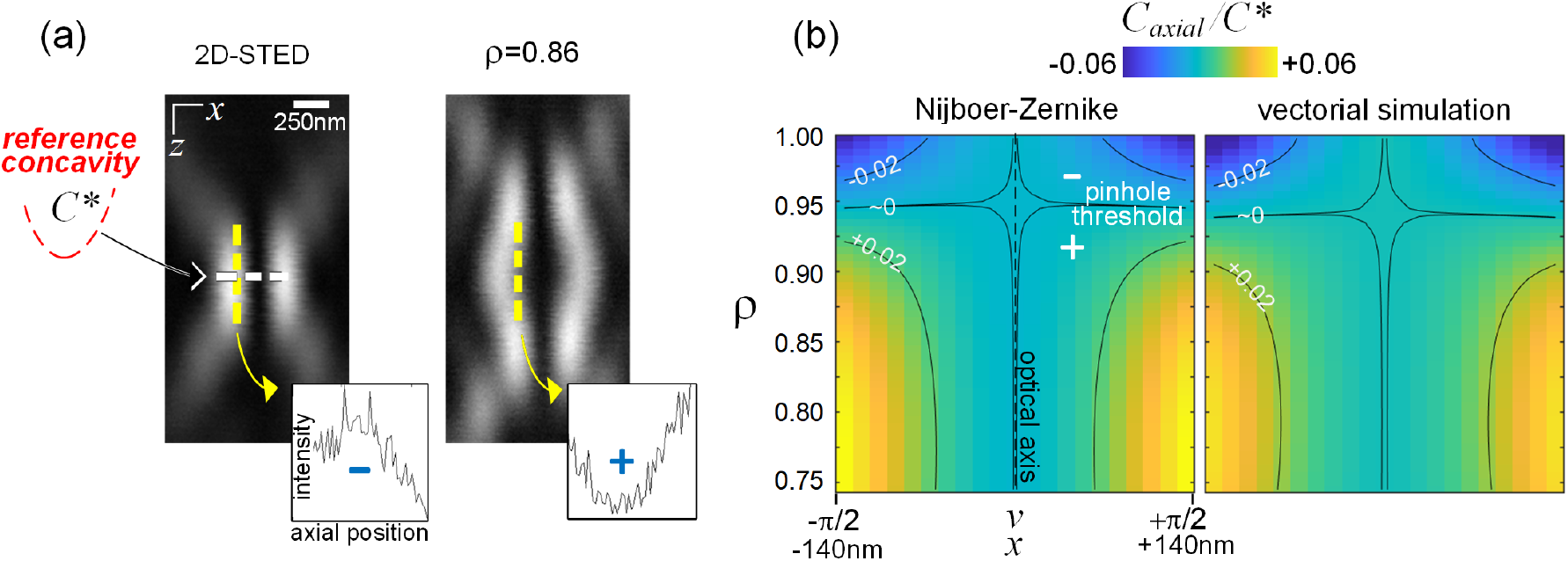
Global pinhole threshold. (a) Guides for measuring axial intensity profile concavities. The concavity of the in-focus lateral profile in 2D-STED is taken as the reference level (*C*\*). (b) Concavity maps as a function of *ρ* and lateral coordinate (x or *v*) showing transition from negative (blue) to positive (yellow) axial confinement at approximately ρ=0.94.

Axial concavity magnitudes (Figure 3a and Eq. S25) lie in the range ±0.05 of the reference value *C* ^*^, which is defined as the maximal concavity in any direction for any depletion mode:

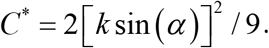

Despite such a comparatively low confinement level, it effectively corresponds to an almost total inversion in relation to the 2D-STED *axial* concavity, which had the wrong sign.

Overall, an increasing contribution of the outer vortex as *ρ* = 1 → 0.8 exchanges some lateral confinement for improved axial confinement, with this flexibility creating the tunable regime of interest (Figure 4).

**Figure 4.**
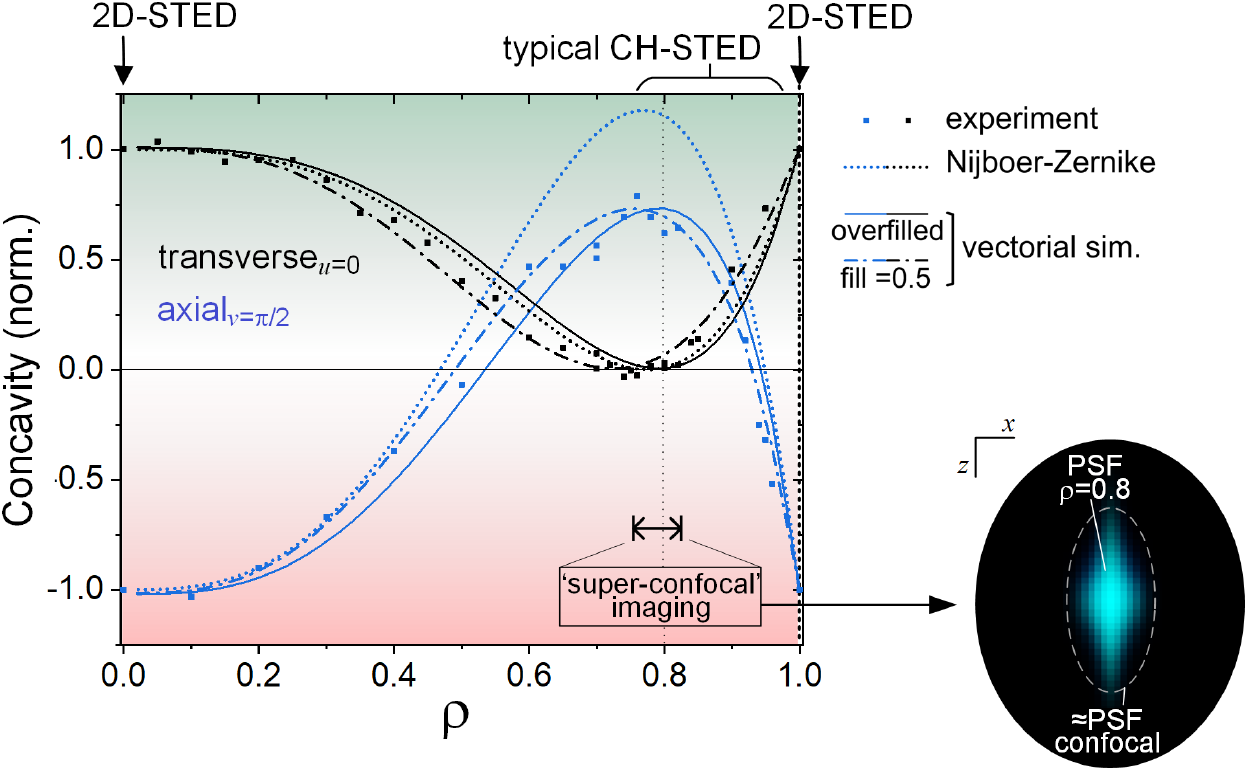
ρ-dependence of lateral and axial confinement. Experiment and theoretical results for lateral confinement (at the focal plane, z=0) and axial confinement (at π/2 (140nm) distance from the optical axis). Vectorial simulation is shown also for a finite waist Gaussian beam. All data is normalized to the absolute values at ρ=1 or 0 (2D-STED). The range 0.8<ρ<1 is most adequate to traverse the full scale of axial and lateral confinement: ρ=1 for full lateral confinement and ρ=0.8 for mostly axial. Inset: Simulated confocal-detection (1 Airy unit) example of a fluorescence effective-PSF at ρ=0.8.

At *ρ ≈*0.8 lateral confinement mostly vanishes (only terms above quartic remain), hinting at a mere diffraction-limited lateral resolution even at high STED saturation. Axial confinement level however peaks at this position, defining an interesting high-signal regime that can be described as super-confocal (Figure 4).

### Optical sectioning across STED modes

Performance of the STED microscope is determined by the impact of depletion on fluorescence, which defines an effective PSF (e-PSF). The question arises as to what extent the observed lateral-axial confinement trends observed in the depletion beam (Figure 4) ultimately lead to enhancements in spatial resolution and optical sectioning in fluorescence imaging. A qualitative assessment using fluorescent samples (Figure 5), namely sub-diffraction beads (0D), microtubule filaments (1D) and a homogeneous plane (2D) (**Methods**) supports the widely held belief that 2D-STED provides the best lateral resolution possible (Figure 5a), as well as the worst sectioning.

**Figure 5.**
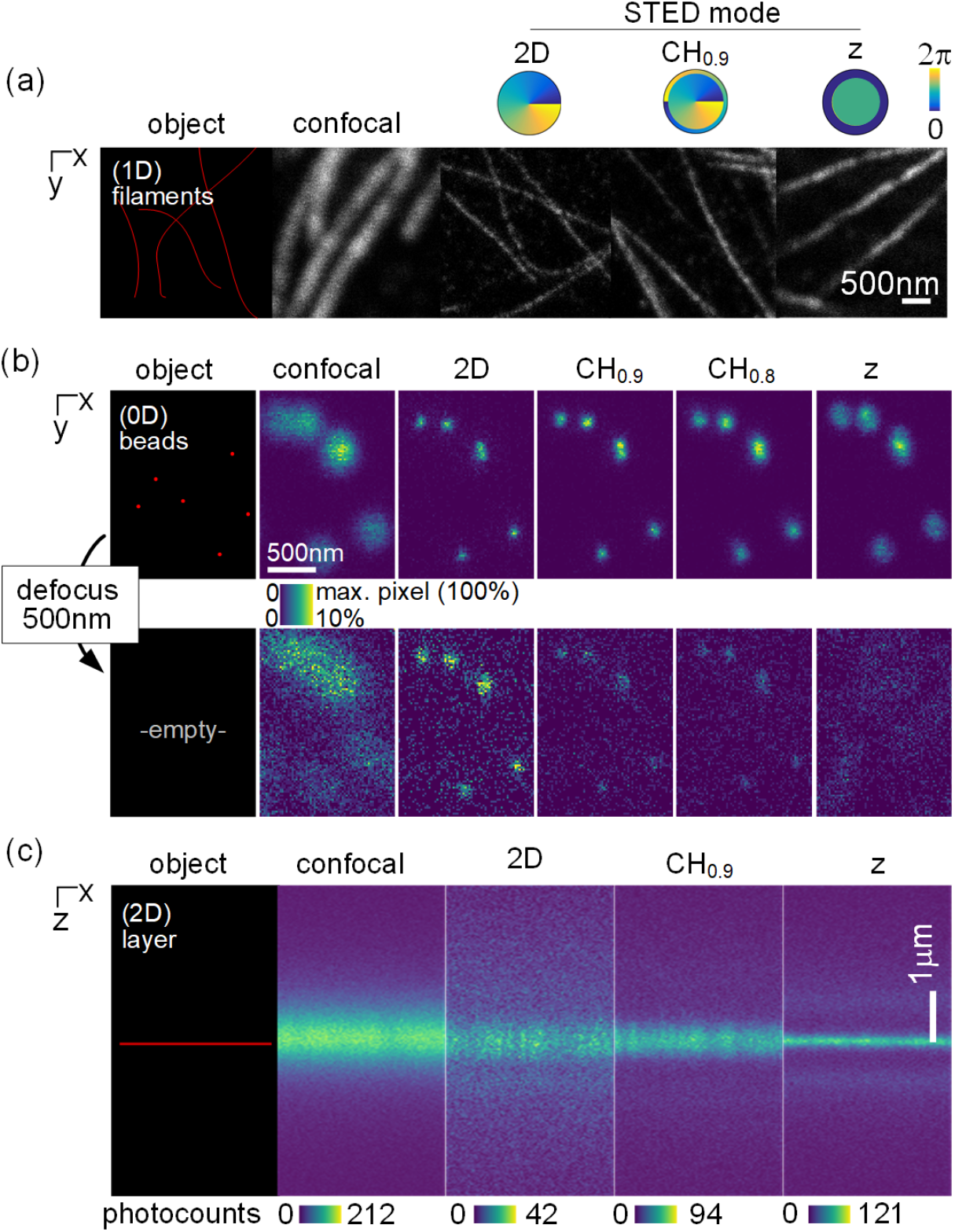
STED imaging of 0D, 1D and 2D objects. (a) Microtubules in Indian muntjac fibroblasts labeled with fluorescent antibody (Abberior STAR Red). (b) xy scanning of 40nm beads). (c) Axial scanning of fluorescent antibody-filled Concanvalin A thin film, forming a quasi-homogeneous planar source. All data using λ_*exc*_=640nm, λ_*STED*_=775nm, λ_*det*_=680nm *n*=1.52, N.A.=1.4, pinhole = 1 Airy unit.

The latter can be clearly observed in the 500nm-defocused images of fluorescent beads (Figure 5b), which retain relatively more signal in the 2D-STED mode than in all other modes. This is a fingerprint of a thick optical slice, or poor optical sectioning. In total contrast, z-STED provides the best sectioning and the worst lateral resolution, clearly revealing a gap between techniques.

Optical slice thickness does not fully characterize optical sectioning. For instance, a section in z-STED is very thin but has ghost replicas at around 800nm from the focal plane (Figure 5c). These arise from the secondary lobes of the excitation PSF not fully encompassed by the z-STED beam geometry. This gives rise to ghost objects in xz scans and less defined background noise in xy scans, which we will later incorporate in the STED metrics.

To allow for a natural comparison among beams that have diverse geometries but carry the same total power, we use a normalized STED power *γ* that refers to the natural common precursor, specifically the peak intensity *I* _*Airy*_ detected using a flat phase mask and a fluorescent point-source. We define *γ* (Figure 6a) as *I* _*Airy*_ normalized by the saturation intensity *I*_*Sat*_ - a property of the fluorescent molecule. Lateral profiles of the effective PSF obtained in a typical confocal setting (overfilled pupil, 1 Airy unit pinhole) are shown in Figure 6b for two values of γ. The vectorial simulation and the Nijboer-Zernike expansion show the advantage of 2D-STED in terms of lateral resolution, with CH0.9-STED, which stands for the CH-STED mode with a bivortex transition radius *ρ* = 0.9, still providing a lateral super-resolution that is far better than that of z-STED at high saturation.

**Figure 6.**
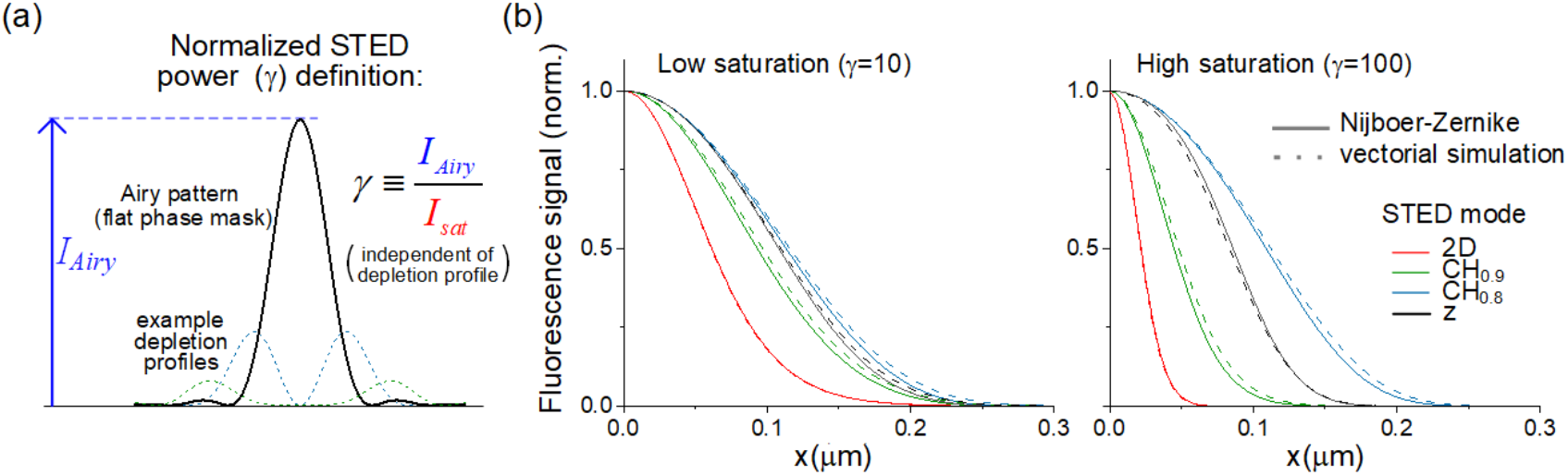
Normalized power and lateral resolution. (a) Definition of the normalized STED power *γ*. (b) Normalized in-focus lateral profiles of fluorescence signal at low and high saturation for various STED modes.

To assess axial performance, we calculated the power at each plane via azimuthal integration (Figure 7a) to obtain optical sectioning curves. The sorting of STED modes in order of performance is mostly reversed (Figure 7b, c) compared to Figure 7a, with the z-STED depletion beam most efficiently providing axial confinement. Both in the Nijboer-Zernike expansion and the vectorial simulation, the well-defined off-focus lobes observed in Figure 5c clearly emerge in z-STED, especially at high *γ*, whilst 2D-STED suffers from delocalized background.

**Figure 7.**
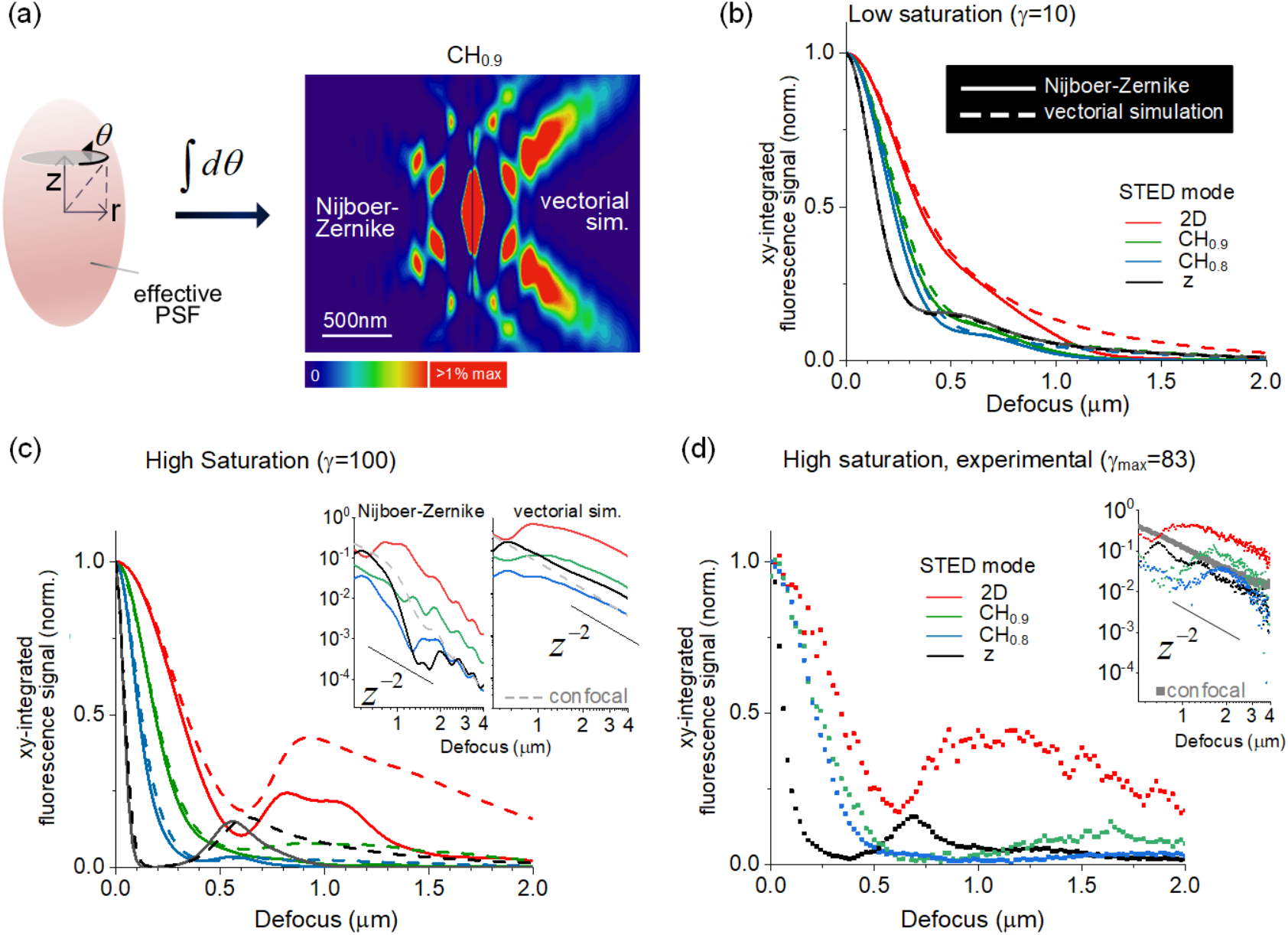
Optical sectioning. (a) The effective PSF rotational integration. (b) Optical sectioning theoretical curves at low saturation and, in (c), at high saturation. (d) Experimental optical sectioning curves at high saturation. Insets in (c) and (d) show curves at large defocus along with the inverse square decay that is expected for all modes at large defocus.

By observing the CH-STED curves’ tails (Figure 7b, c), it seems clear that the CH-STED curves are not mere interpolators between 2D- and z-STED. They are so in terms of lateral and axial confinement, but not in terms of background elimination, where CH-STED seems optimal. To test this premise experimentally, we measured sectioning curves using a sufficiently dense ensemble of point sources distributed in 2D (the fluorescent planes as in Figure 5c), equivalent to an azimuthal integration of the PSF. To obtain the data (Figure 7d) we used the maximum depletion laser power available in our setup, against an Abberior STAR-Red fluorescent layer, yielding a normalized power of *γ* = 83. Experimental sectioning curves show slice thickness variations and off-focus lobes/tails that are broadly consistent with theory both in position and amplitude and confirm background elimination superiority of CH-STED compared to both conventional STED modes. We will make these assessments quantitative in the next section.

The Nijboer-Zernike results only diverge from the full vectorial simulation and the experimental data at large defocus values with high depletion power while using a weak-sectioning mode, namely the CH0.9. Essentially, truncation of the Nijboer-Zernike expansion leads to signal underestimation, but this only becomes significant for positions that are far from the optical axis (due to the ‘long’ azimuthal integration) while still inside the ‘geometric beam cone’, so that actual signal is still relatively high. These two conditions are met only for points far from the focal plane (Figure 7a).

At large defocus, all experimental curves display higher signal plateaus than the simulations, as corruption by noise increases. Still, the inverse-square law asymptotic behavior that is expected from geometrical optics is observed in both the experimental and vectorial data, while the Nijboer-Zernike prediction – a (finite) expansion rooted at the PSF center - falls off faster (see the insets in Figures 7c, d).

### Resolution, sectioning, and signal space

To compare the STED modes, we consider three figures of merit: lateral resolution, optical section thickness and an optical section quality factor, Q, which measures the signal-to-background conditions (Figure 8). We define Q as the fraction of detected signal emitted from inside the nominal optical slice, so that a distinction can be made between curves that yield a similar slice thickness (e.g. FWHM) while their off-focus behavior may actually reveal large differences, such as the presence of secondary lobes or a long tail. One example is the axial performance of z-STED at γ=10 (black curves in Figure 7b) compared to CH0.8 at γ=100 (blue curves in Figure 7c), which have comparable slice thickness but will have different Q values because of dissimilar off-focus behavior.

**Figure 8.**
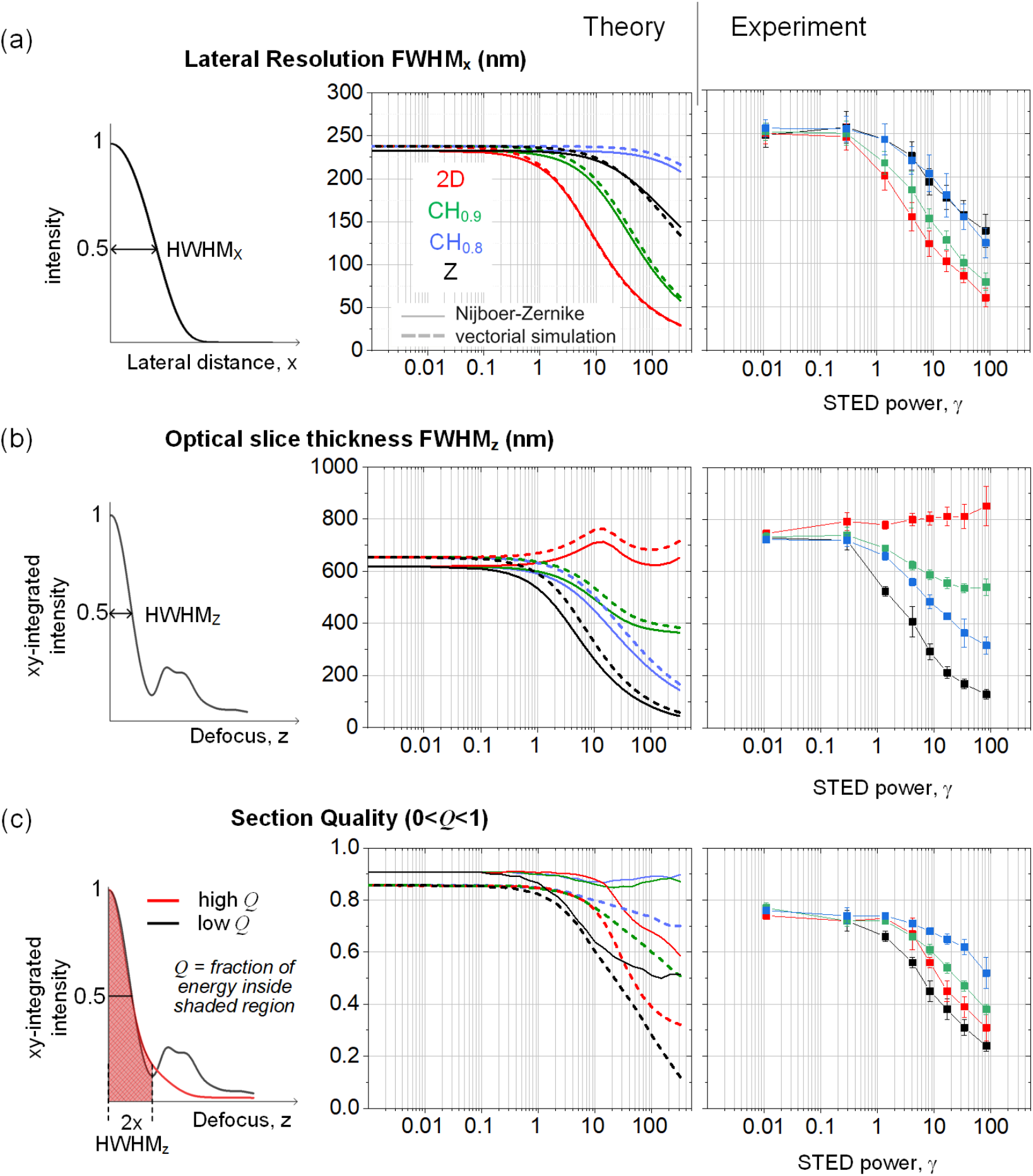
Saturation level and saturation geometry dependencies. Criteria for (a) Lateral resolution, (b) Optical slice thickness and (c) Section Quality (Q) are depicted (left column) and the experimental and theoretical results are presented as a function of normalized depletion power. Error bars represent *s*.*d*.

The two main control parameters in the CH-STED microscope are the usual depletion laser power, characterized by *γ*, that alters the saturation *level*, and the bivortex transition radius *ρ* (and topological charge, so that z-STED is included in the comparison), which alters the saturation *geometry*. For clarity, we avoid the continuous spectrum of CH-STED bivortices and limit the analysis to the set of four representative situations: 2D-STED, z-STED and two CH geometries (Figure 8). CH-STED modes interpolate between the two other modes regarding lateral resolution and slice thickness (Figure 8a, b), emulating 3D-STED with only one beam. What is however more striking is that, while bridging the two standard STED modes, CH-STED does so with better a better signal-to-background ratio (Q) than either mode (Figure 8c). This is also evident in the mammalian cell imaging of Figure 9a, and is particularly noticeable at very high saturation (Figure 9b), when loss of SNR becomes the more critical, actually exacerbating the inherent disadvantages and exposing any particular defects of the 2D-STED beam.

**Figure 9.**
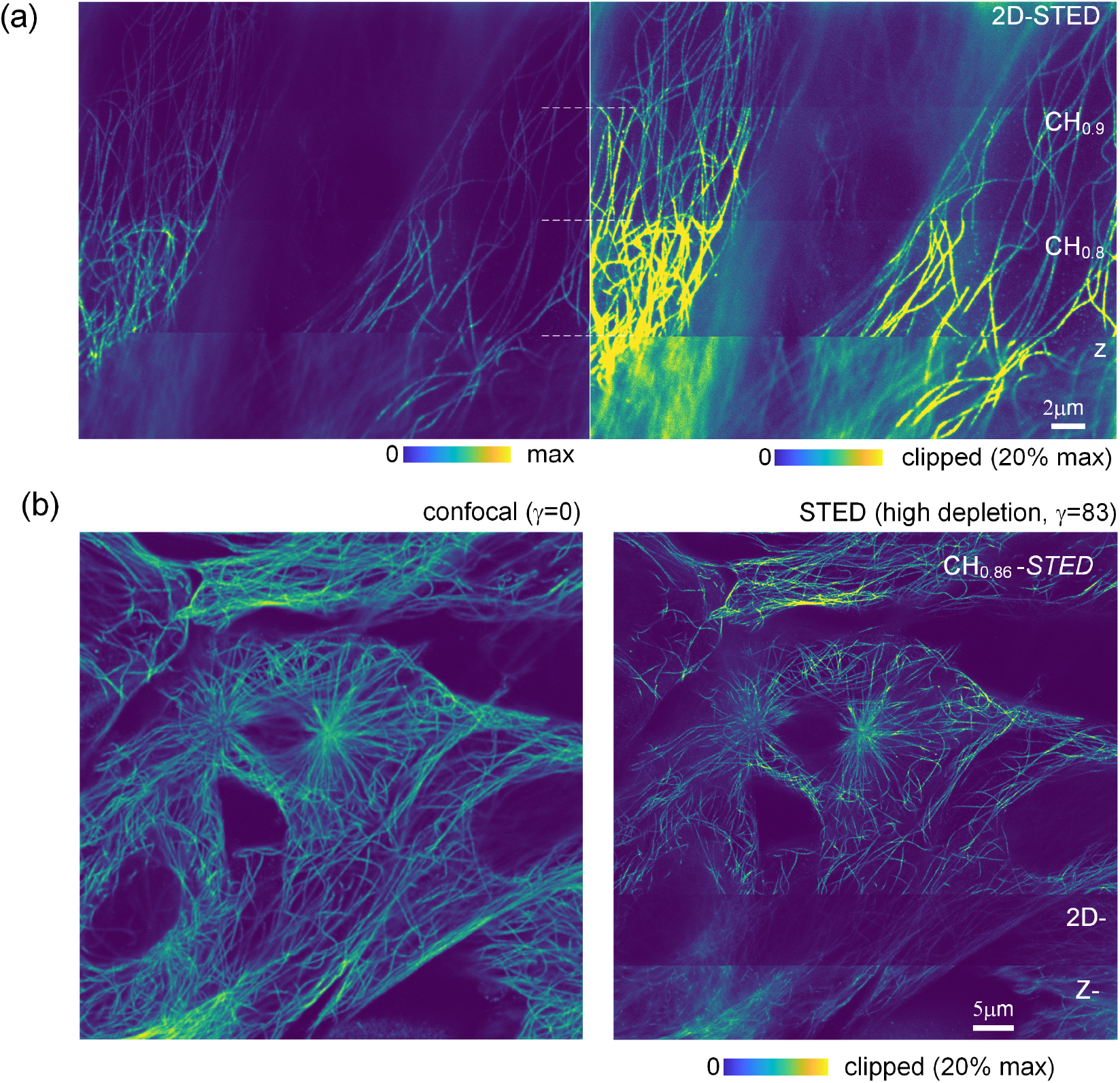
STED cell imaging: Microtubules inside Indian Muntjac fibroblast cells labeled with a (large) fluorescent antibody. a) STED xy scan (slow axis is vertical) with on-the-fly switching of the SLM’s phase mask that modulates the depletion laser. b) High-power STED imaging (γ=83) so that 2D-STED sensitivity to instrumental deficiencies of the STED beam is actually exacerbated.

We note that the estimation of contrast, as measured by the factor Q, is the most sensitive to signal derived from points distant from the focal plane, presenting a greater challenge to the Nijboer-Zernike expansion, whose predictions over-estimate performance relative to simulation and experiment.

Figure 10 clearly shows that CH-STED fills a previously inaccessible area of the parametric space (Figure 1c). Here each separate contour corresponds to a given value of the bivortex vortex transition radius, *ρ*, while progress along a contour displays the evolution as the saturation parameter, γ, varies. The experimental data for the CH0.8 mode departs significantly from theory under the overfilling approximation (blue dots in Figure 10), which leads to an overestimation of the power fraction transmitted across the outer vortex of the phase plate. To first order this can be accounted for in the model by correcting nominal *ρ* with a larger effective *ρ*_*eff*_ adjusted so that the ratio of power in the inner and outer ring for incident plane waves is equal to that of the truncated Gaussian beam at the nominal value of *ρ*. Given such positive offset towards *ρ*_*eff*_, performance tends towards better lateral resolution than expected for the nominal *ρ*. Opposing this bias, the effectively lower numerical aperture provided by the Gaussian beam trivially leads to a loss in lateral resolution.

**Figure 10.**
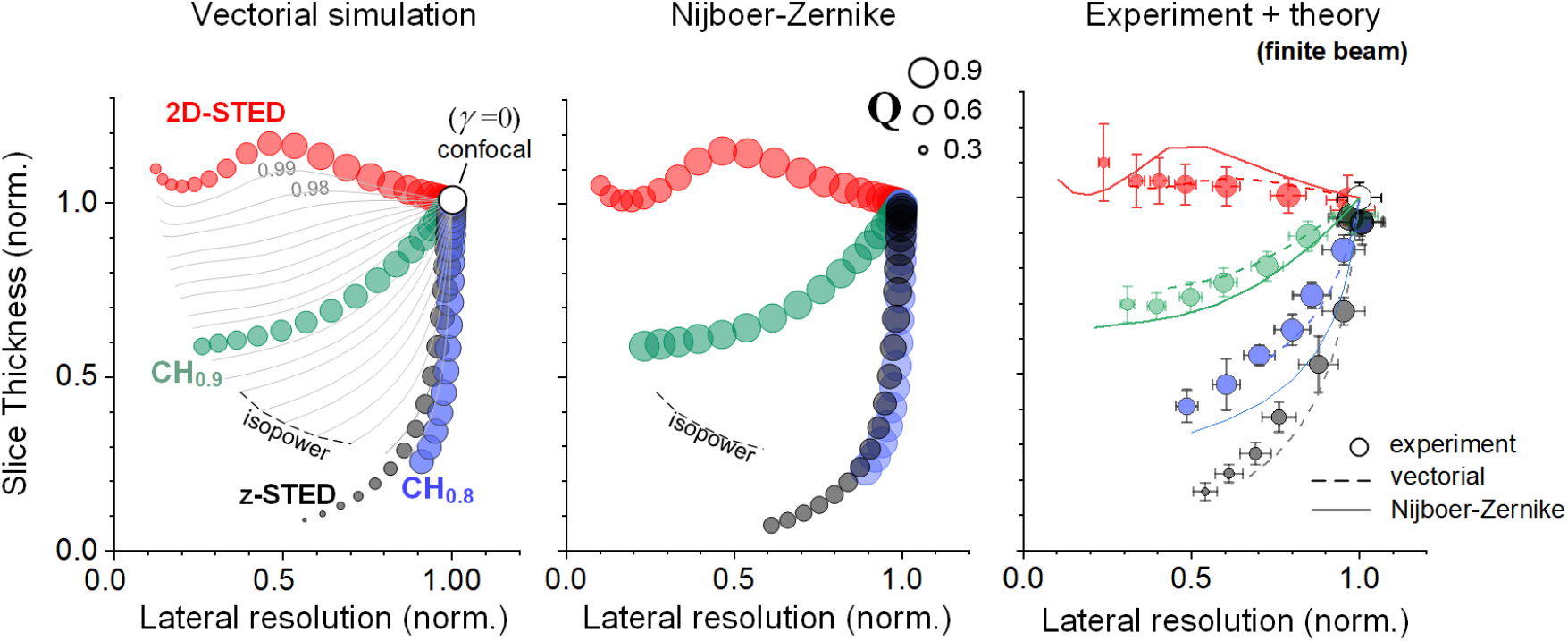
Filling the performance gap. Performance of the various depletion beam parameters displayed in terms of 3 quality metrics: lateral resolution, optical slice thickness, and sectioning quality, Q. The latter is proportional to the diameter of the circle (scale in the middle graph). In all graphs, the two Cartesian coordinates are normalized to the confocal value γ=0 (for absolute values, refer to Figure 8). The experimental data is shown along with the vectorial and Nijboer-Zernike predictions when the finite width of the depletion beam is used. Error bars represent *s*.*d*.

These trends were analyzed using the full vectorial simulation of a truncated Gaussian beam and the usual Nijboer-Zernike expansion with a recalculated *ρ*_*eff*_. The two calculations are shown along with the experimental data, showing the expected bias (Figure 10, right) in relation to the plane-wave assumption (Figure 10, left and center).

## Discussion

For almost 20 years, the 2D-STED/ z-STED beams have defined the complete toolbox in STED microscopes, with end-users eventually having the possibility to adjust the balance between the two depletion ‘strengths’ by independently varying the incident powers of the two superimposed beams. Subsequent advances have pertained mostly to dimensions other than PSF geometry, such as the identification of the PSF center via fluorescence time-signatures during depletion^37-40^, removal of background with multi-acquisition protocols^41, 42^, enhanced probes and labeling strategies ^43, 44^, intelligent illumination schemes^45^ among other developments aimed at either improving imaging or decreasing photo-toxicity (for reviews, see for example ^12, 23, 46^). Optimizations focused on PSF geometry have been reported but were focused on improving lateral resolution (even if with increased lateral lobes) ^47, 48^. To our knowledge, proposals prior to the bivortex beam did not account for higher, less resolution-centric metrics, such as one including signal-to-background.

A prime motivation to look for alternatives to the vortex was the understanding that its design is scale-less, an advantage in robustness, but barring any tunability. The top-hat, on its hand, has two design parameters, the disc’s phase step and radius, but these are not tunable. On the contrary, both parameters have to be very precisely set. The bivortex has the same parameters, but one can freely set them, allowing for a smooth trade-off between lateral resolution and better axial confinement. This opens up a much larger space that can be finely sampled to optimize various aspects of performance. Generally, the quality of images obtained from more densely packed samples will benefit from this flexibility.

Given its outright inefficiency at the optical axis, the benefits of the bivortex bottle beam are not immediately apparent. At first glance, the axial node appears to be a structural ‘defect’ that compromises axial confinement. The nodal line would indeed be compromising if the purpose was, for example, to trap a small (low-index) particle, which might indeed escape through the well. Likewise, the fluorescence profile of the CH-STED e-PSF along the optical axis is exactly equal to that of a confocal or a 2D-STED microscope. This seems to imply that axial resolution is not improved, but this is only (tendentially) true for the highest transverse modulation frequencies. Actually, the level of confocality (or optical sectioning) of an imaging system is determined by the axial resolution at the lower edge of the transverse frequencies (*k*_*x, y*_ = 0, the OTF’s axial profile), and this is given by the unbounded lateral integration of the intensity PSF, as a function of defocus. This DC-background rejection is what more strongly exposes the sectioning advantage of CH-STED (e.g. Figure 8b), although the entire low-frequency domain will actually show improved axial resolution.

The Nijboer-Zernike expansion used throughout the study proved to be computationally efficient and to perform well in comparison with the full vectorial diffraction simulation in predicting resolution and sectioning. It thus provides an efficient means for the analysis of microscope performance that can be useful in developing pre-optimization procedures, even in high-NA optical systems.

## Supplemental Material

### Focal field using the Nijboer-Zernike expansion

Here, we outline a heuristic pseudo-paraxial description of the different STED depletion beam geometries. While there are more accurate methods available ^3, 49^, we feel that the simplicity of the approach taken here based on the original Nijboer-Zernike formalism for describing weak aberrations allows one to obtain a good qualitative description quickly. In particular, it enables one to readily explore curvature metrics described in the main text.

#### a) Three dimensional light distribution near the focal plane

We describe the electric field amplitude distribution near the focal plane of an ideal lens within the scalar Debye approximation of the Huygens-Fresnel diffraction integral. In this approach, the objective is essentially a phase converter, transforming the planes waves incident on the entrance pupil of an ideal infinity–corrected objective into spherical waves converging on the geometric focus. Starting from equation (4) of section 8.8 in Born and Wolf’s seminal work^31^, an incident field *U*_*in*_ is transformed by an ideal objective lens with focal length *f* according to the Debye-Wolf integral,

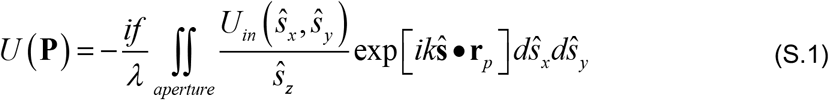

**P** represents a general point in the vicinity of the on-axis focal plane with coordinates **r**_*p*_ = (*r* cosψ, *r* sinψ, *z*). Here, the unit vector 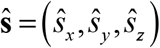 describes the direction of a ray from an arbitrary location on the spherical exit pupil towards the geometric focus. Transverse coordinates of 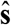 describe positions within the aperture of the lens’ exit pupil. The wavenumber k is that of the incident light in the medium. Given the exit pupil’s spherical nature it is convenient to write 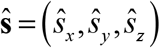 in terms of spherical coordinates, 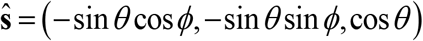. We assume the incident field to be planes waves of amplitude, A, modified by a phase mask M that will determine the type of STED beam created. In this case, the Debye-Wolf integral becomes,

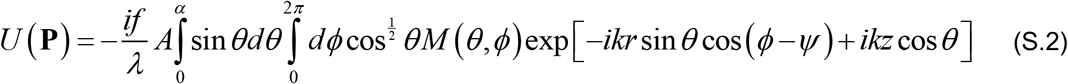

The factor of cos^1/2^ *θ* arises from the apodization factor of an objective that satisfies the Abbe sine condition. The limiting angle of the objective’s exit pupil is *α* related to the objective’s numerical aperture, *NA* = *n* sin *α* through the medium’s refractive index, *n*. At this point it is convenient to introduce scaled axial and radial coordinates (*u, v*)in the image space,

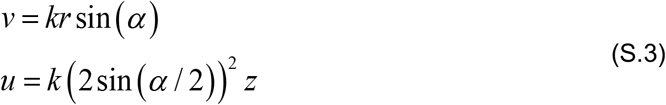

We note that this choice for the scaled axial position, *u*, advocated by Sheppard and Matthews^33^ differs from that adopted by Born and Wolf, namely *u* = *k* sin^2^ *α*, although they are equivalent in the paraxial limit. Using these scaled coordinates the argument of the integrals becomes,

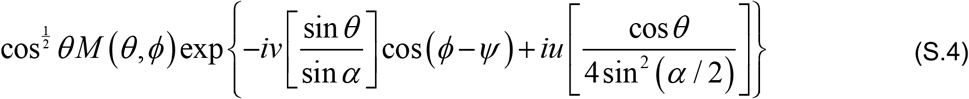

To make further progress, we make a change of variables sin*θ* = *σ* sin *α* and invoke the pseudo-paraxial approximation^33^ for high numerical apertures,

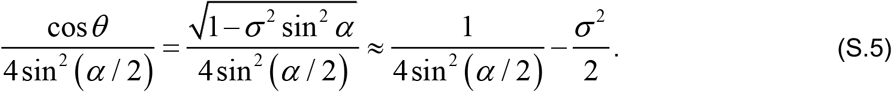

This approximation has the virtue of agreeing with the full expression at the two limits of the integral *σ* = 0 and *σ* = 1. In contrast, the conventional paraxial approximation expands the square root in the above equation about *σ* = 0, giving 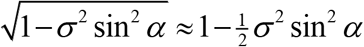. In the paraxial limit, *α* ≪ 1 this yields the correct curvature on-axis, but diverges more quickly from the full expression as *α* increases.

Putting this altogether, we arrive at the following approximation for the scalar field of a phase modulated plane wave in the vicinity of the focal region of a high-numerical,

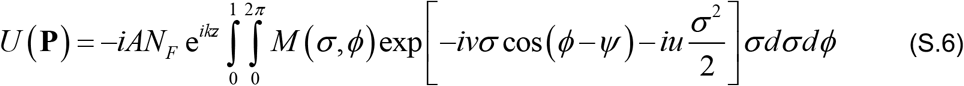

Here, *N*_*F*_ = *f* sin^2^ *α* / λ is the Fresnel number for the objective with an angular aperture *α*. Under typical conditions (a vacuum wavelength of 775nm, a 60x objective with a numerical aperture of 1.4 and a refractive index of 1.52) the Fresnel number is of order 3650. For simplicity, we have also chosen to neglect the apodization weighting factor [1− *σ* ^2^ sin^2^ *α*]^−1/ 4^ appropriate to an objective that obeys the Abbe sine condition. If included this factor would place a somewhat higher weight on rays with larger values of *σ* and would tend to create an asymmetry between the intensity at equal distances before and after the focal plane. By neglecting it we effectively force the intensity patterns to be symmetric about the focal plane.

We now consider four main cases for the phase-mask *M* (*σ, θ*). For incident plane waves, the phase mask can be written as

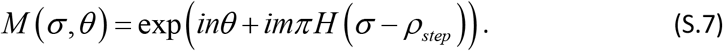

with *H* representing the Heaviside step function.

The simplest case is that of a circular pupil with no effective phase mask for which *n* = *m* = 0 .

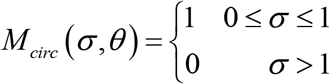

The integral over *ϕ* in Eq. (S.1) gives a zeroth order Bessel function, leaving

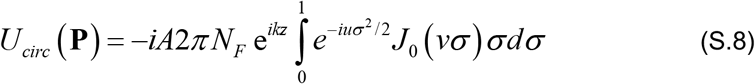

We use the on-axis in focal plane amplitude of this case as a normalization for the various STED profiles. On-axis in the focal plane,

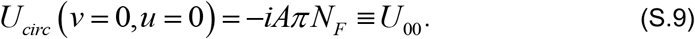

The intensity at the peak of the Airy spot *I* _*Airy*_, is proportional to |*U*_00_ |^2^.

For a general position near the focal spot,

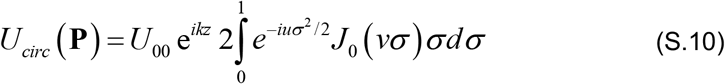

To make progress, we use a substitution originally proposed by B.R.A. Nijboer^50^ in developing his description of diffraction in the presence of minor aberrations. It involves expanding the exponential under the integral in terms of Legendre polynomials, *P*_*s*_ (*x*) which can be related to a subset of the radial Zernike polynomials, 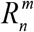 :

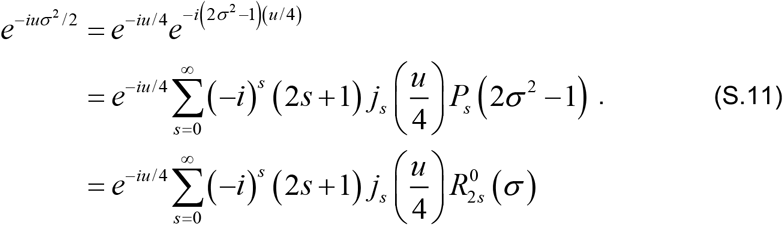

Here 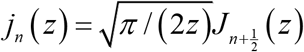 are the spherical Bessel functions of the first kind.

The chief advantage of this expansion is that one can make use of the remarkable Nijboer’s result

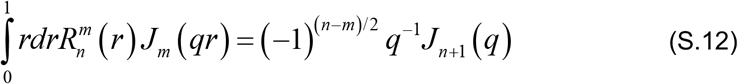

to carry out the integral over *ρ*. This gives

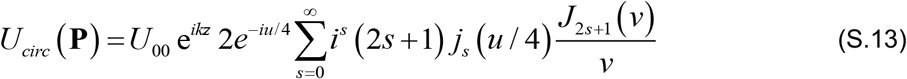

Conveniently this series seems to converge very quickly and the first four terms are sufficient to obtain and intensity profile fairly accurate for (*u, v*) ranging from 0 to ±10*π*. When compared to the full vector simulation for circularly polarized incident light on a high numerical aperture system (NA = 1.4, n=1.52, and λ= 775 nm) the first four terms of the Nijboer-Zernike expansion are accurate to within 5% percent. We note that the Debye approximation makes the intensity profiles symmetric about both the axial and lateral axes, while the full vectorial simulation leads to slightly higher intensities for positive z. This asymmetry is accentuated for the STED depletion beam intensity profiles because of the interference inherent in their construction.

**Figure S1.**
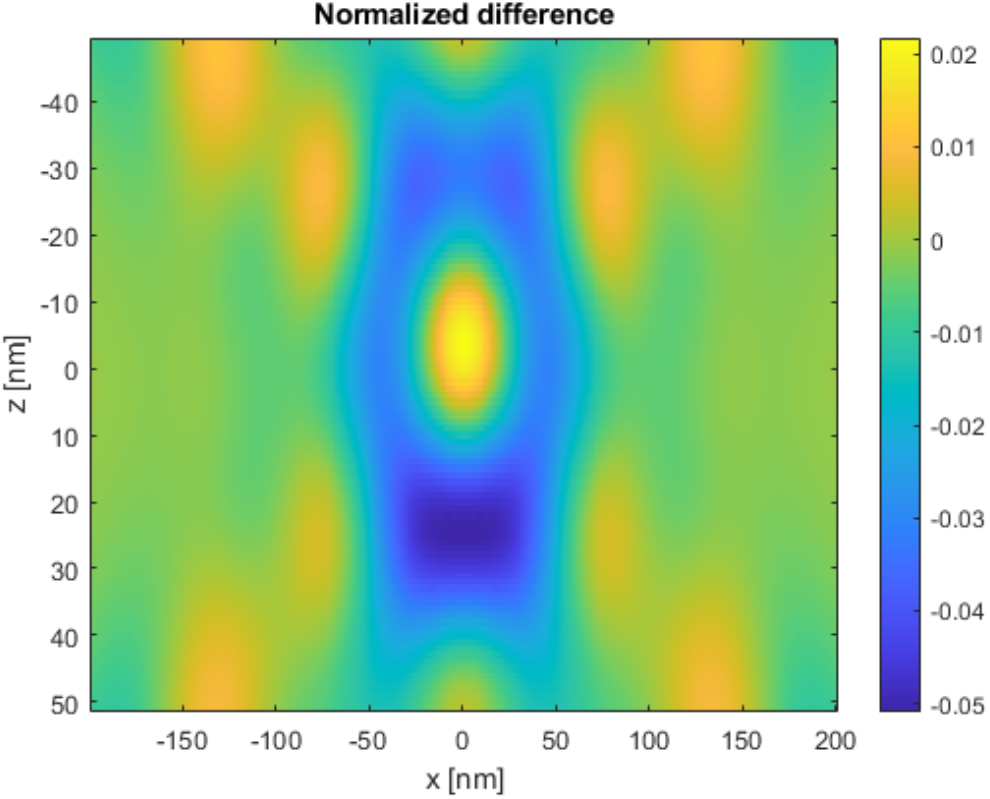
Nijboer-Zernike versus vectorial simulation. Difference between the intensity calculated from the first four terms of the scalar pseudo-paraxial Nijboer-Zernike expansion (Eq. S10) and the full vectorial simulation for circularly polarized incident light at 775 nm with a flat phase mask focused by an ideal objective with NA = 1.4 with a refractive index of 1.52 in image space.

The next simplest case is the z-STED profile with a phase mask that is the difference between an inner circular aperture and an outer ring. For incident plane waves the division of the pupil into two equal areas occurs when 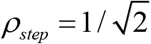 such that

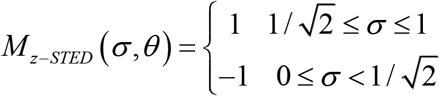

This gives immediately

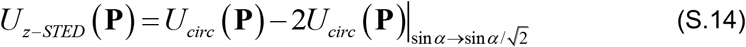

Using our scaled coordinates, when sin*α → ρ* sin*α*,

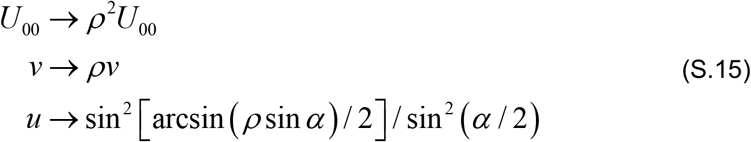

For the single Vortex (2D-STED) the phase mask is simply

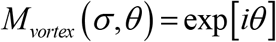

and in this case the angular integral results in a fist order Bessel function:

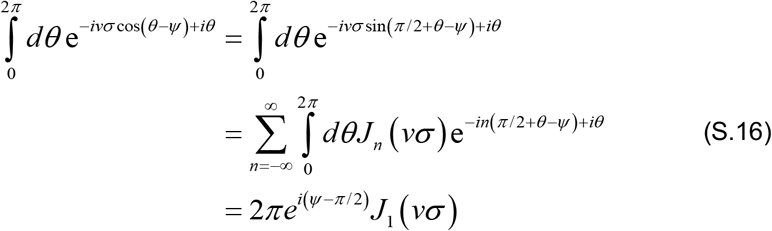

Combining this result with the Nijboer expansion of Eq. S.11 gives

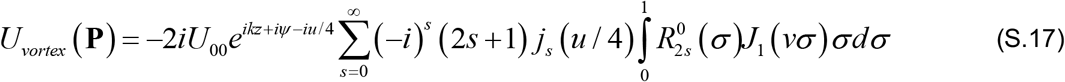

Unfortunately, now the order of the Bessel function differs from the upper index of the radial Zernike polynomial, so Eq. S.12 is no longer valid. Nevertheless, one can express the relevant integral using generalized hypergeometric functions:

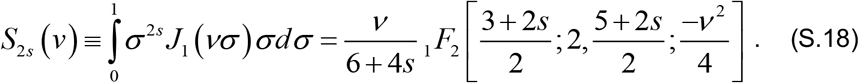

Here the generalized hypergeometric functions are 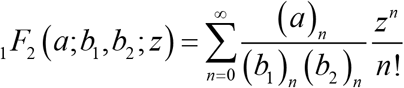, with 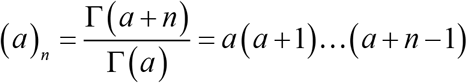 being the Pochhammer symbol.

Making use of the explicit form for the first few radial Zernike polynomials

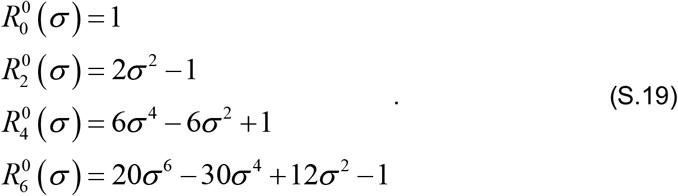

The end result for the first four terms in the series expansion is

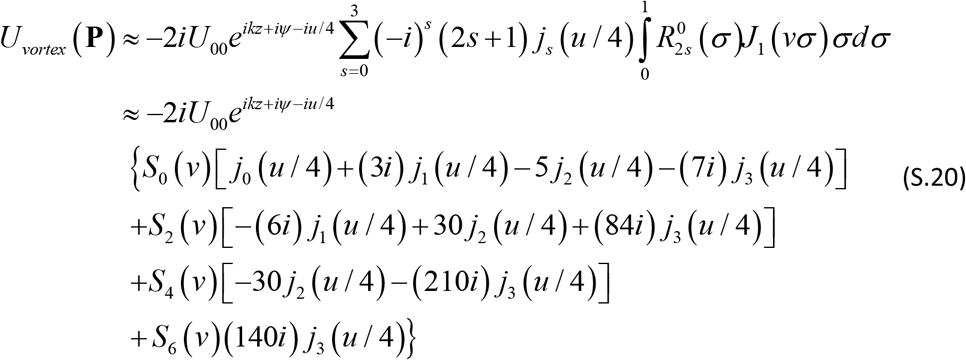

Finally, we consider the bivortex (CH-STED), a coherent combination of a (deliberately) decalibrated z-STED mask and a single vortex. We denote by *ρ* the normalized radius at which the vortex switches sign, i.e.

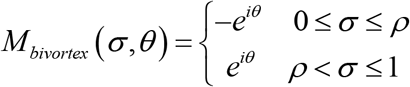

Then in analogy with the z-STED case,

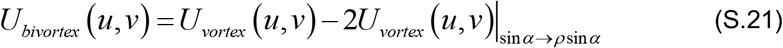

#### b) Estimates of the curvature metric

the normalized radius at which One advantage of the Nijboer-Zernike expansions is that one can readily carry out Taylor expansions and estimate the curvature metric referred in the main text as a measure of the confinement capacity of each STED mode. The maximum curvature is realized for 2D-STED on axis (*u* = 0)in the transverse direction. Given that the leading term in the Taylor expansion of the spherical Bessel functions about the origin scale as *j*_*n*_ (*u*) ∼ *u*^*n*^, only the *j*_0_ (*u* / 4)term in the expansion of Eq. S.20 contributes to the 2D-STED intensity profile in the focal plane,

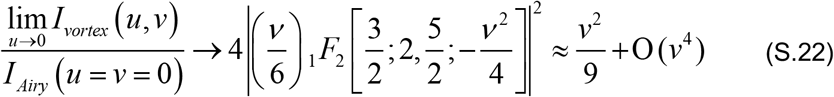

leading to an estimate of the maximum curvature in scaled radial coordinates of

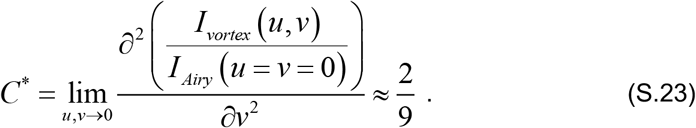

In physical units, i.e. going from v back to r, the reference curvature is *C*^*^ = 2 [*k* sin (*α*)7]^2^ / 9.

In contrast, the axial curvature of the 2D-STED at an arbitrary scaled transverse coordinate, *v* is given by coefficient of the quadratic term in the scaled axial coordinate, *u*^2^ when expanding the field about the focal plane. We find that

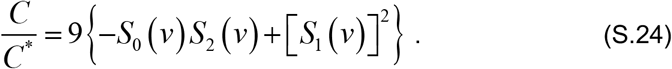

As shown in Figure S2, the axial curvature for the 2D-STED beam is negative well beyond the strong axial lobes which extend to approximately 3π/4, reflecting the total lack of axial confinement of the 2D-STED depletion beam throughout the excited volume.

**Figure S2.**
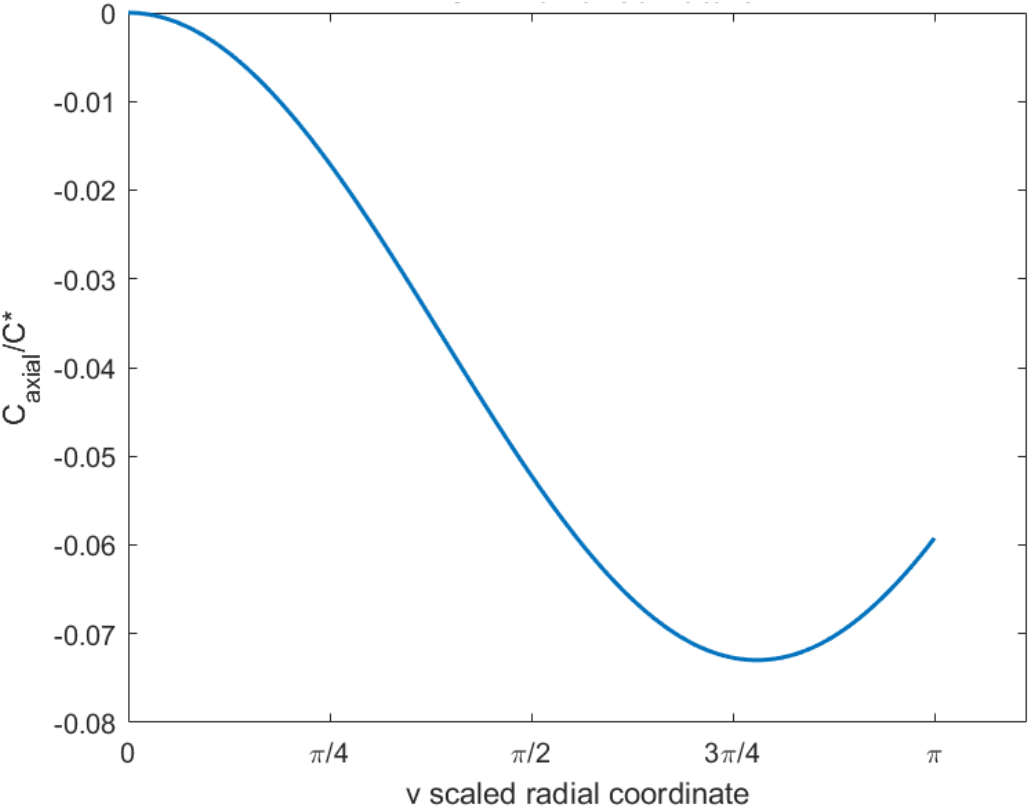
Lateral dependence of the axial curvature in 2D-STED. The axial curvature metric for the 2D-STED beam within the truncated Nijboer-Zernike expansion as a function of the scaled radial coordinate v. Up to v = π the axial curvature is negative indicating a lack of axial confinement. At v = π/2 the normalized curvature is C_axial_/C* ≈ -0.053.

For the CH-STED beam the situation is different, there is a large of *ρ* values that provoke a positive curvature metric for the CH-STED beam. Directly on axis, at *v* = 0, the axial curvature vanishes. Using Eqs. S.20 and S.21 for the Nijboer Zernike expansion to calculate the CH-STED intensity and Taylor-expanding the result about the focal plane (u=0) we find that up to terms quadratic in u, the intensity of the CH-STED beam near the focal plane is given by

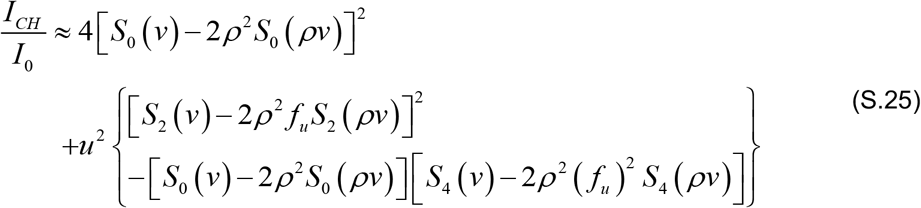

Here *f*_*u*_ is the change in the normalized axial coordinate, u due to the reduced solid angle when the aperture is reduced by a multiplicative factor of *ρ*, i.e.

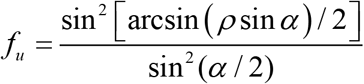

At this level of approximation, the curvature will switch sign when the coefficient of the *u*^2^ term sums to zero. The resulting scaled curvature metric is shown in Figure S.3 for the case when v =π *π*/2. For this case the curvature metric is positive for *ρ* between (0.46 and 0.95) with a maximum value of 0.062 at *ρ* = 0.77.

**Figure S3.**
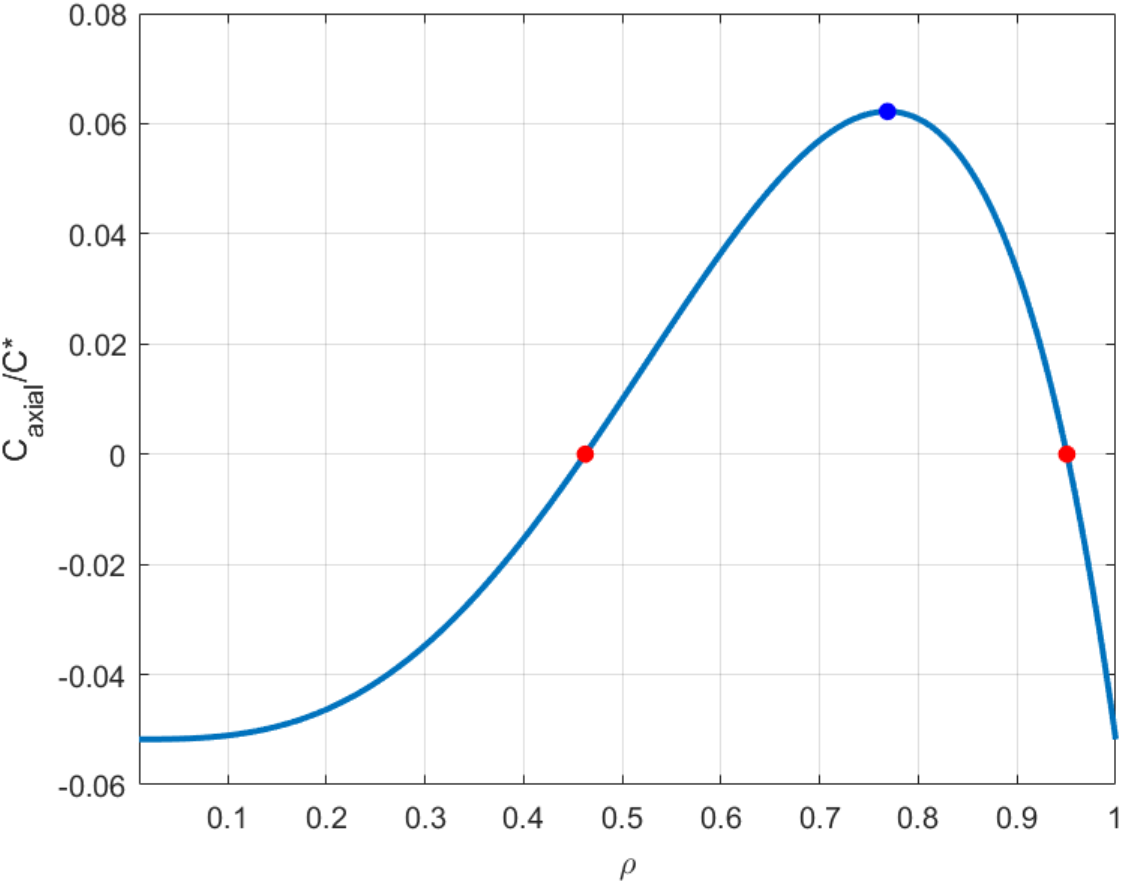
ρ-dependence of the axial curvature in CH-STED. The axial curvature metric for the CH-STED beam as estimated from the truncated Nijboer-Zernike expansion as a function of the scaled transition radius *ρ* when *v* = π/2 close to the axial position corresponding to radius at which the depletion beam’s strong intensity lobes reach a maximum. The axial curvature switches sign at the points *ρ* = 0.46 and *ρ* = 0.95. The maximum is approximately 0.85 times the absolute value of the most negative curvature metric estimated for the 2D-STED beam.

The transition points at which the axial curvature switches sign vary only slightly with the scaled radial coordinate *v*, as shown in Figure S.4.

**Figure S4.**
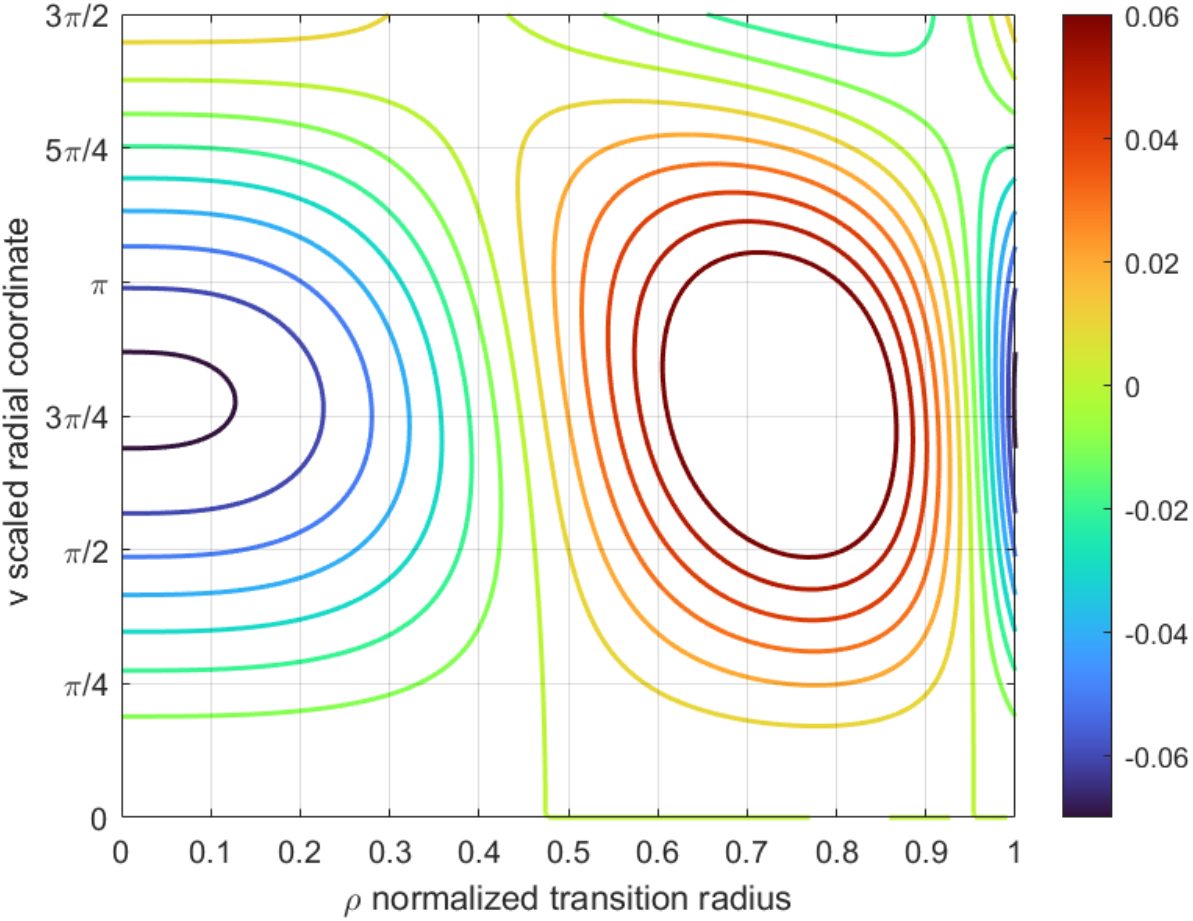
Positional and r-dependence of axial curvature in CH-STED. Variation of C_axial_/C* as a function of the scaled radial coordinate v and the normalized transition aperture radius, *ρ* There is a wide range of *ρ* values that lead to a positive (confining) axial curvature metric, extending from just below 0.5 up to 0.95 as indicated by the yellow-green contours. In practice, one can obtain enhanced axial confinement over a wide radial extent for *ρ* close to 0.9. Here, the simulation was carried out for an objective with a numerical aperture of 1.4, an image space refractive index of 1.52 and an incident (vacuum) wavelength of 775 nm.

## Methods

### Indian muntjac cells (IM)

IM fibroblasts, immortalized with human hTERT (pBabe puro hTERT, kind gift from Jerry W Shay), were grown in Minimum Essential Media (MEM) (Gibco, Life Technologies), supplemented with 10% FBS (Gibco, Life Technologies), at 37 ºC in humidified conditions with 5% CO2. IM fibroblast cells were seeded on fibronectin-coated coverslips 2 days before the experiment. After fixation with 4 % Paraformaldehyde and 0.2 % Glutaraldehyde (Electron Microscopy Sciences) diluted in cytoskeleton buffer (274 mM NaCl, 10 mM KCl, 2.2 mM Na2HPO4, 0.8 mM KH2PO4, 4 mM EGTA, 4 mM MgCl2, 10 mM Pipes, 10 mM glucose, pH 6.1). After fixation, the cells were quenched with a 0.1% solution of Sodium Borohydrate (Sigma-Aldrich) in PBS and extracted using PBS-0,5% TritonX (Sigma-Aldrich) diluted in cytoskeleton buffer. The coverslips were incubated with the primary antibody [anti-tyrosinated tubulin 1:150 (MCA77G, Bio-Rad)] in blocking solution (10% FBS with 0.05% Tween 20 diluted in cytoskeleton buffer overnight at 4ºC. After washing with PBS-0.05%Tween 20, coverslips were incubated with the secondary antibody [Abberior STAR RED 1:100 (Abberior Instruments)] for 1h at room temperature. DAPI (4’6’-Diamidino-2-phenylindole, Sigma Aldrich) was then added for 5 minutes in PBS-0.05% Tween 1:50,000. Coverslips were washed in PBS and sealed on glass slides using mounting medium (20 nM Tris pH 8, 0.5 N-propyl gallate, 90% glycerol).

### Fluorescent antibody-filled Concanavalin A film

A solution of 0.5 mg/ml Concanavalin A was mixed with secondary antibody [Abberior STAR RED 1:100 (Abberior Instruments)]. Plasma cleaned coverslips were coated with this solution and incubated for 1h at room temperature. After washing with PBS, the coated coverslips were sealed on glass slides using mounting medium.

### Fluorescent beads

A drop of fluorescent beads suspension (TetraSpeck™ microspheres, Invitrogen) was spread on the surface of a glass slide until dry. The sample was then covered with mounting media and a glass coverslip.

### Image display

All acquired images are shown ‘raw’, subjected to linear histogram adjustment, always starting at 0 photo-counts.

### *ρ* calibration

Gold bead scattering (detected without a pinhole) at the depletion laser wavelength (775nm) was used to calibrate the scale of the phase masks. Flat-field-corrected masks are written on a spatial light modulator (SLM) that is imaged approximately onto the back focal plane of the microscope objective lens (Nikon Lambda VC 60x 1.4NA). To calibrate the SLM effective dimension we first inscribed a vortex (helical phase) pattern because it is devoid of a radial scale. As such, centering the vortex mask at the optical axis (judged by symmetry of the doughnut pattern) is sufficient to create the typical 2D-STED depletion pattern, regardless of the scale calibration status. Concavity of the parabolic intensity profile near the 2D-STED doughnut dip was measured and defined as a reference value *C*_*ref*_. With a bivortex template, a *ρ*_*in*_ value was coarsely determined to be inside the pupil radius, as judged by any definite change in the pattern morphology, specifically an axial spreading detected with xz scanning. Concavity values were then precisely determined in a range around *ρ*_*in*_. Concavity values are approximated by (1− 2*ρ* ^3^)^2^ and thus have an approximately linear dependence with *ρ* in a short range below the (not exactly known) pupil radius. We used the range 0.9-0.95 of the (not exactly known) pupil radius to perform a linear fit. An extrapolation of the linear fit to *C*_*ref*_, the intersection defining radius of the effective pupil.

### Fill Factor Determination

One practical problem with z-STED is that the intensity at the doughnut dip reaches zero only for the specific radius, *ρ*_*top*−*hat*_, that equalizes the effective contributions of each mask ring to avoid de-excitation of the sample on-axis at focus. However, in this specific context empirical knowledge of the top-hat radius that is required to achieve z-STED (^*ρ*^*top*−*hat* =0.62) provides a very practical means to infer the filling ratio of the pupil by the incident Gaussian beam. Using the nominal settings of the relevant components (NA=1.4, n=1.518, λ=775nm) we looped through the simulation for different filling ratios to determine the value that minimized the dip intensity. Filling ratio was determined to be 0.47. Note that the observation that the lateral confinement curve in Figure 4 does not reach a minimum at around *ρ*_*bivortex*_ ≅0.79 but at a lower value already hinted that the fill factor was finite.

**Figure S5.**
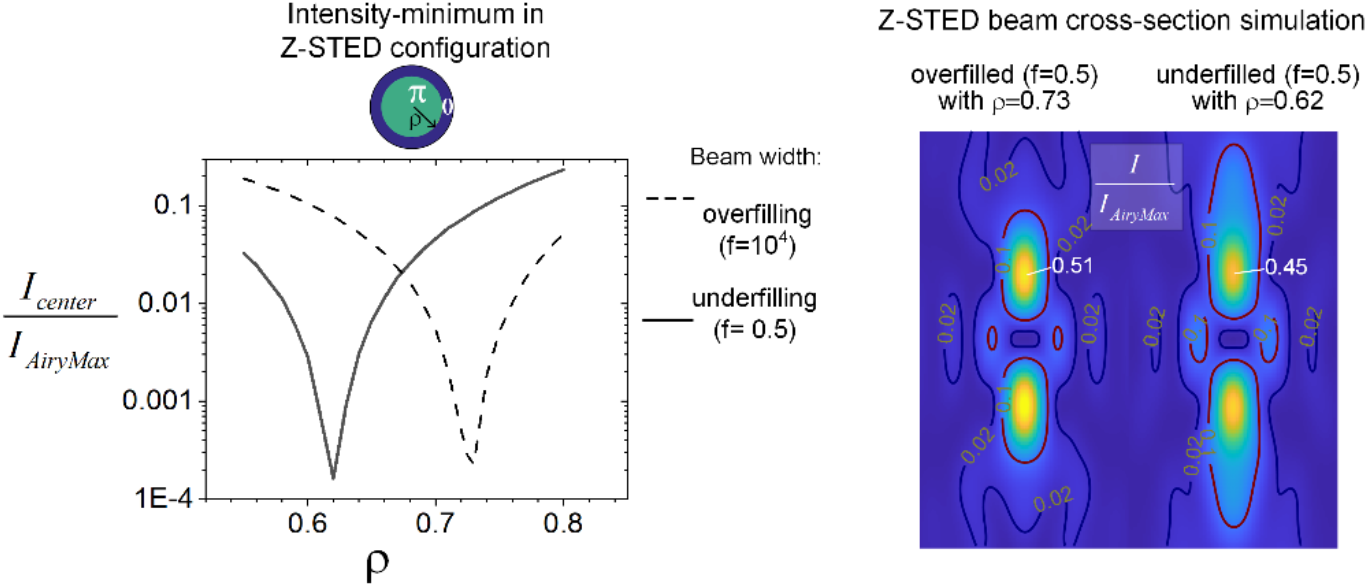
Fill factor determination. A finite fill factor attenuates contribution from the outer parts of the phase mask, forcing the z-STED top-hat radius to be lower than would be for a plane wave so that. the intensity-zero at the center is retrieved.

### Normalized power (γ) calibration

STED imaging using a Gaussian depletion beam instead of the doughnut beam was used to determine the depletion power that caused the signal of STAR-Red-decorated sample to decrease to 1/e relative to confocal level.

### Vectorial simulations

All vectorial simulations were performed using the Focus Field Calculator Matlab package^29^.

## Acknowledgements

The authors thank Helder Maiato for providing conditions for the development of the work and to members of his group, to Jorge Ferreira and José Carlos Gomes for discussions. A.J.P work is done under a position granted by the program Estímulo ao Emprego Científico (CEEC) from the portuguese Fundação para a Ciência e a Tecnologia (FCT). This work was funded by the project: DOI 10.54499/EXPL/NAN-OPT/0476/2021.

